# Fate mapping melanoma persister cells through regression and into recurrent disease in adult zebrafish

**DOI:** 10.1101/2022.03.17.484741

**Authors:** Jana Travnickova, Sarah Muise, Sonia Wojciechowska, Alessandro Brombin, Zhiqiang Zeng, Adelaide I.J. Young, Cameron Wyatt, E. Elizabeth Patton

## Abstract

Melanoma heterogeneity and plasticity underlie therapy resistance. Some tumour cells possess innate resistance, while others reprogramme during drug exposure and survive to form persister cells, a source of potential cancer cells for recurrent disease. Tracing individual melanoma cell populations through tumour regression and into recurrent disease remains largely unexplored, in part, because complex animal models are required for live imaging of cell populations over time. Here, we apply tamoxifen-inducible *cre^ERt2^/loxP* lineage tracing to a zebrafish model of MITF-dependent melanoma regression and recurrence to image and trace cell populations *in vivo* through disease stages. Using this strategy, we show that melanoma persister cells at the minimal residual disease site originate from the primary tumour. Next, we fate mapped rare MITF-independent persister cells and demonstrate that these cells directly contribute to progressive disease. Multiplex immunohistochemistry confirmed MITF-independent persister cells give rise to Mitfa+ cells in recurrent disease. Taken together, our work reveals a direct contribution of persister cell populations to recurrent disease, and provides a resource for lineage tracing methodology in adult zebrafish cancer models.

**Summary statement:** We fate map melanoma cells from the primary tumour into a persister cell state and show that persister cells directly contribute to recurrent disease.

## INTRODUCTION

Melanoma, a deadly cancer of pigment producing melanocytes, ranks amongst the highest for genetic and transcriptional heterogeneity (Rambow et al., 2019; Travnickova and Patton, 2021). Therapy resistance remains a major challenge for patients with melanoma, with partial or short-term responses characteristic of targeted therapy, eventually resulting in tumour relapse (Patton et al., 2021; Shen et al., 2020b). This resistance can be intrinsic, in which pre-existing primary tumour cell states directly confer resistance, or acquired, whereby melanoma cells develop resistance upon drug exposure either by acquiring new genetic mutations or adapting their transcriptional state (Marin-Bejar et al., 2021; Marine et al., 2020; Shen *et al.,* 2020b). These resistance mechanisms likely occur concurrently, and contribute to the high heterogeneity within melanoma and the persister cell states at the minimal residual disease (MRD) site.

Recently, using single cell RNA sequencing (scRNA-seq), we and others have demonstrated high transcriptional heterogeneity and distinct cell states in the primary tumour and MRD, including states with low-to-no expression of pigmentation lineage markers such as melanocyte inducing transcription factor (MITF)-independent populations (Baron et al., 2020; Ennen et al., 2015; Gerber et al., 2017; Rambow et al., 2018b; Tirosh et al., 2016; Travnickova et al., 2019). These studies have identified multiple melanoma cell states that have been proposed to drive tumour recurrence, while others are of unknown contribution to disease progression (Travnickova and Patton, 2021). While revealing new concepts about melanoma transcriptional cell states, these studies reflect only a single or few time points, and are thereby limited in what can be understood about the behaviour of individual cell populations over time.

This limitation is particularly critical in the context of cell plasticity in cancer models. Advances in imaging technologies coupled with lineage tracing methods in zebrafish models now enable the fate mapping of cell populations over time (Mosimann et al., 2011; Pan et al., 2013). The power of an inducible lineage tracing system is illustrated by advances in understanding of organ development and tissue homeostasis in zebrafish (Carney and Mosimann, 2018; Thunemann et al., 2017). For example, lineage tracing experiments using the general neural crest marker *sox10* or the recently established melanocyte stem cell marker *tfap2b* provided proof of the existence of multipotent melanocyte precursors during early embryonic development and their contribution to adult zebrafish pigment cell patterning (Brombin et al., 2022; Singh et al., 2016). Similarly, a conditional *cre/loxP* recombination system demonstrated the hierarchy of neuroepithelial progenitors and the functional heterogeneity of neural stem cells in the vertebrate adult brain using neural lineage specific markers (Galant et al., 2016; Than-Trong et al., 2020). Furthermore, multispectral lineage tracing revealed the mechanism behind myotome generation (Nguyen et al., 2017). The regenerative capacities of zebrafish combined with an inducible lineage tracing system have been instrumental in understanding the cell origin and lineage restrictions in regenerated organs such as lesioned heart, fins or spinal cord (Briona et al., 2015; Jopling et al., 2010; Tornini et al., 2017). Recently, multicolour tracing has been combined with mosaic mutagenesis using CRISPR-Cas9 in a novel technique called TWISTR (tissue editing with inducible stem cell tagging via recombination) to show that the fitness of mutant clones is controlled by resistance to inflammation (Avagyan et al., 2021). Despite these advances, the application of tamoxifen to control the temporal and spatial activation of Cre (Cre^ERt2^) in adult zebrafish cancer models has been hampered by technical challenges, such as lack of established protocols and tamoxifen toxicity.

Here, we adapt and optimise the inducible *ubi:Switch* lineage tracing system (Mosimann *et al.,* 2011) for use in an adult zebrafish cancer model to fate map cells through melanoma growth, regression and recurrence. For the first time, we directly capture melanoma cell switching from MITF-independent persister cells to Mitfa positive cells to contribute to recurrent disease *in vivo.* Our work supports the concept that targeting persister cells will be critical to delay or prevent recurrent disease.

## RESULTS

### Conditional tamoxifen induced fluorophore switch in adult zebrafish melanoma

We have previously developed a conditional MITF-dependent zebrafish melanoma model (*Tg*(*mitfa:BRAF^V600E^*);*mitfa^vc7^;tp53^M214K^*), in which melanoma regresses and recurs concurrently with the changes in MITF activity controlled by temperature (Travnickova *et al.,* 2019). In this model, MITF activity is controlled by a temperature sensitive splicing mutation in the *mitfa* gene (*mitfa^vc7;^* zebrafish orthologue *mitfa* is expressed in the body melanocytes) (Johnson et al., 2011; Zeng et al., 2015). At the lower permissive temperature, MITF activity is on and promotes tumourigenesis, while at the higher temperature *mitfa* RNA is still expressed but is not spliced correctly and thereby MITF protein levels and activity are abolished. Turning off MITF activity results in tumour regression with remaining persister cells at the MRD site (Travnickova *et al.,* 2019).

To trace melanoma cells through disease states, we first set out to establish an inducible *cre/loxP* system on the MITF-dependent melanoma background. To this end, we expressed *cre^ERt2^* from the *mitfa* promoter to generate a transgenic *Tg(mitfa:cre^ERt2^)* line and crossed this with a *ubi:Switch* reporter line (Mosimann *et al.,* 2011). This experimental design would enable us to induce a green-to-red (GFP-to-mCherry) permanent fluorophore switch in *mitfa:cre^ERt2^* expressing melanoma cell populations at a defined time point in melanoma disease progression (**Fig. 1A, B**).

**Figure 1:**
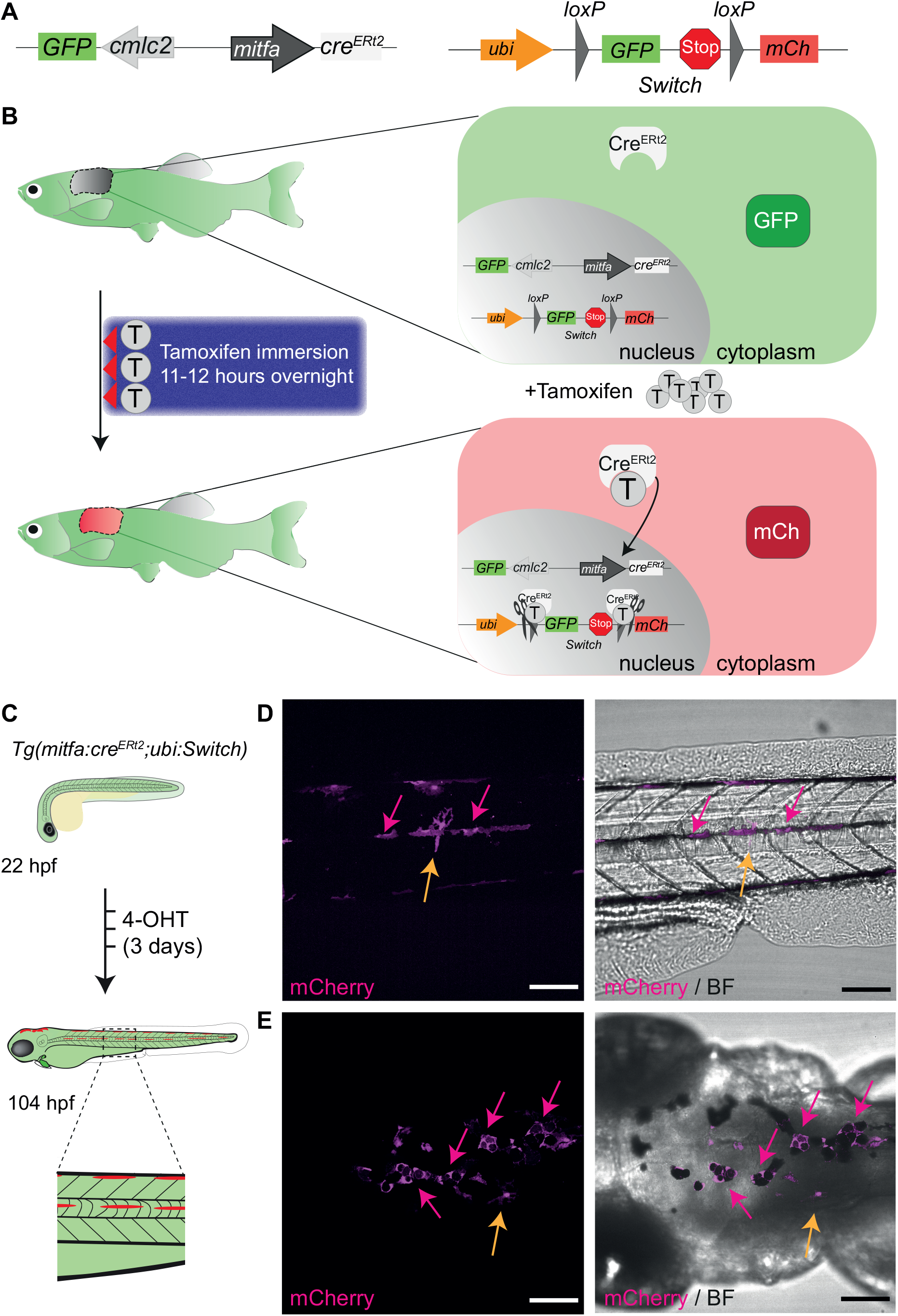
A tamoxifen inducible *mitfa:cre^ERt2^/loxP* system in zebrafish labels melanocytes. **A.** A schematic of *mitfa:cre^ERt2^* (left) and *ubi:Switch* (right) (Mosimann *et al.,* 2011) constructs to enable a green-to-red switch in *mitfa* promoter expressing cells after tamoxifen treatment. The *cardiac myosin light chain (cmlc)* 2 promoter drives *GFP* expression and facilitates screening for the *cre^ERt2^* line based on GFP+ heart myocardium. mCh: mCherry. **B.** A schematic of the concept behind the colour switching *cre^ERt2^/loxP* system that allows green-to-red switch after tamoxifen treatment in cells expressing the *mitfa* promoter driving *cre^ERt2^.* Left: A three-night-course treatment with tamoxifen for 11-12 hours by immersion in the dark with a drug-free period during the day. Right: The principle of colour switching upon tamoxifen treatment as a result of Cre^ERt2^ trans-localisation to the nucleus, causing excision of *GFP* and a stop codon, enabling mCherry expression in recombined cells. **C.** An illustration of the expected green-to-red fluorophore switch in the zebrafish embryo upon three consecutive daily treatments with 20μM 4-hydroxytamoxifen (4-OHT). Enlarged view of dashed line rectangle shows “switched” melanocytes in the embryonic stripes. **D.-E.** Lateral (**D**) and dorsal (**E**) views of a zebrafish larva after 4-OHT treatment at 104 hpf (4.5 dpf) validates *mitfa:cre^ERt2^* specificity to melanocyte lineage. Standard deviation intensity (STD) projection of a confocal z-stack (left; mCherry) and single z-plane acquisition (right, brightfield (BF)) at 6 hours after the end of 4-OHT treatment. mCherry expressing melanocytes are visible and express black melanin pigment (pink arrows). Two unpigmented, star shaped cells expressing mCherry are also visible and may represent xanthophore progenitors (orange arrows). N = 6 fish for each view, scale bar = 100 μm

To validate the specificity of the *mitfa:cre^ERt2^* construct for *mitfa* expressing cells, we first performed the fluorophore switch during early embryonic development using 4-hydroxytamoxifen treatment (4-OHT) (**Fig. 1C**). During embryogenesis and in regeneration, Mitfa is required for the generation of melanocytes and *mitfa* expression marks melanoblasts and progenitors of additional pigment cells, including the yellow xanthophores (Brombin *et al.,* 2022; Parichy et al., 2000). Upon 4-OHT treatment (three consecutive daily treatments of 20 μM), we could detect mCherry expressing cells at 4.5 days post fertilisation (dpf) along the lateral stipe and on top of the head, in both cases co-localised with pigment (**Fig. 1 D-E,** pink arrows). Some of the mCherry positive cells (**Fig. 1D-E**, yellow arrow) that did not co-localise with pigmented cells likely correspond to pigment cell progenitors (Brombin *et al.,* 2022; Parichy *et al.,* 2000). This indicates that *mitfa:cre^ERt2^* construct specifically labels *mitfa* expressing cells.

We found that 4 μM tamoxifen treatment of adult zebrafish by immersion for three consecutive nights (11-hour-treatment, 13-hour-recovery) was successful for *cre^ERt2^/loxP* recombination without toxicity (**Fig 1B**, **Fig. 2A**). We chose tamoxifen over 4-OHT for its increased solubility in dimethylsulfoxide (DMSO) which is well tolerated by adult zebrafish. The treatment was completed overnight to align with the natural light-dark cycle of the fish and to prevent phototoxicity of tamoxifen (Wang et al., 2009). By three days post tamoxifen treatment course, we could detect mCherry in the primary tumour, but not in DMSO-treated controls (**Fig.2B**, **Fig. S1A**). Using confocal microscopy, we validated the presence of individual clusters of mCherry+ melanoma cells in tamoxifen treated tumours only (**Fig. 2C-D**). Immunostaining of melanoma sections with an mCherry antibody showed that mCherry+ cells were present both at the surface of the tumour and in the invading melanoma cells along muscle fibres thus confirming the efficiency of this protocol to fate map cells throughout the body of the melanoma tumour (**Fig. 2E, F**).

**Figure 2:**
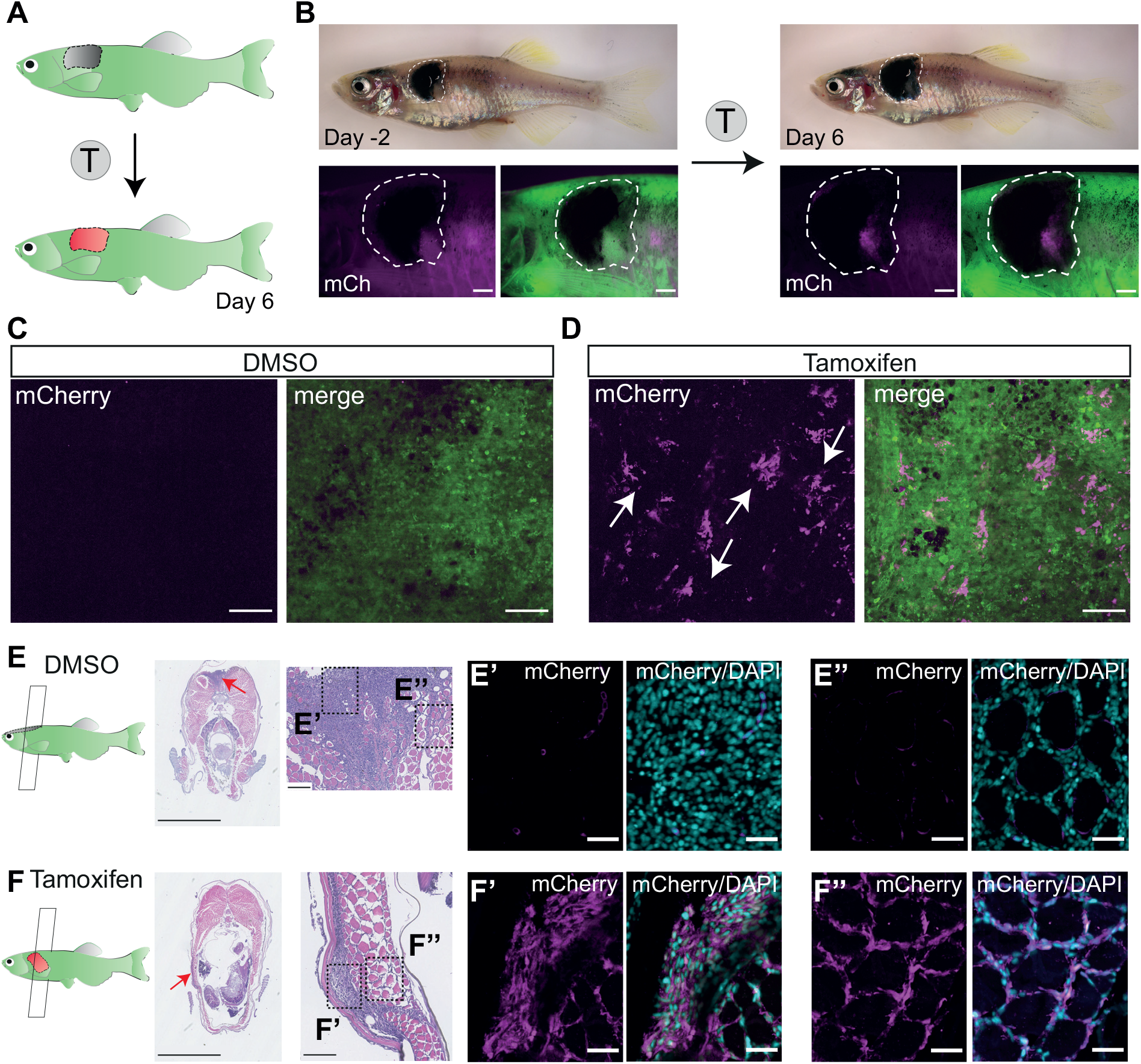
Successful fluorophore switch in adult zebrafish primary melanoma after tamoxifen treatment. **A.** An illustration of the expected fluorophore switch in the primary melanoma tumour upon tamoxifen (T) treatment and imaged 6 days after the start of the treatment. **B.** An adult zebrafish with a fully developed primary melanoma tumour (delineated with white dashed line) 2 days before the tamoxifen (T) treatment course (left) and 6 days after the start of the treatment (right) showing *de novo* expression of mCherry protein in pigmented trunk tumour after tamoxifen treatment (4 μM). N = 2 fish, Scale bar = 1 mm. **C.-D.** STD projections of confocal z-stacks of DMSO (0.04 %) treated fish (**C**, left mCherry, right merged with GFP) and tamoxifen treated fish (**D**, left mCherry, right merge) show small clusters of mCherry expressing cells in the tamoxifen treated group only (white arrows). N = 2 fish, scale bar = 100 μm. **E.-F.** Paraformaldehyde fixed paraffin embedded (FFPE) transverse sections of primary melanoma fish (left) stained with haematoxylin and eosin, with zoomed images of the tumour (red arrows pointing to the location of the zoomed images). N = 2 fish, scale bars = 2.5 mm and 100 μm respectively. Dashed boxes indicate area of section shown in **E’-F’** and **E”-F”**. **E’.-F’.** Immunofluorescence staining of bulk tumour with a mCherry antibody (magenta), counterstained with the nuclear marker DAPI (cyan), shows mCherry signal in tamoxifen treated samples only (**F’**) compared to DMSO control (**E’**). N = 2 fish, scale bar = 25 μm. **E’’.-F’’.** Immunofluorescence staining of invasive tumour show successful fluorophore switching and mCherry expression in the tamoxifen treated group only (**F’’**) compared to DMSO control (**E’’**). N = 2 fish, scale bar = 25 μm. See also **Figure S1A**.

Having established the system in a single colour switch reporter line, we wanted to test if our method was applicable to the multicolour system of fate mapping. To this end, we crossed the *Tg(mitfa:cre^ERt2^)* melanoma prone fish with the *ubi:zebrabow* transgenic line (Pan *et al.,* 2013) to evaluate permanent colour changes in *cre^ERt2^* expressing cells from default red (RFP) to a stochastic combinatorial expression of three fluorophores upon tamoxifen treatment: cyan (CFP), yellow (YFP) and red (RFP) (**Fig. 3A**). Indeed, similar to *ubi:Switch* line, we could detect *de novo* fluorescent signal by three days post tamoxifen treatment compared to DMSO controls, mainly in the YFP channel, and at 34 days post treatment, we could detect all three fluorophores within the primary melanoma (**Fig. 3B, Fig. S1B**). Using confocal microscopy, we validated the presence of multicolour labelling (CFP, YFP and RFP), which allows distinction of individual cells across the labelled tissue (**Fig. 3C, Fig. S1C**).

**Figure 3:**
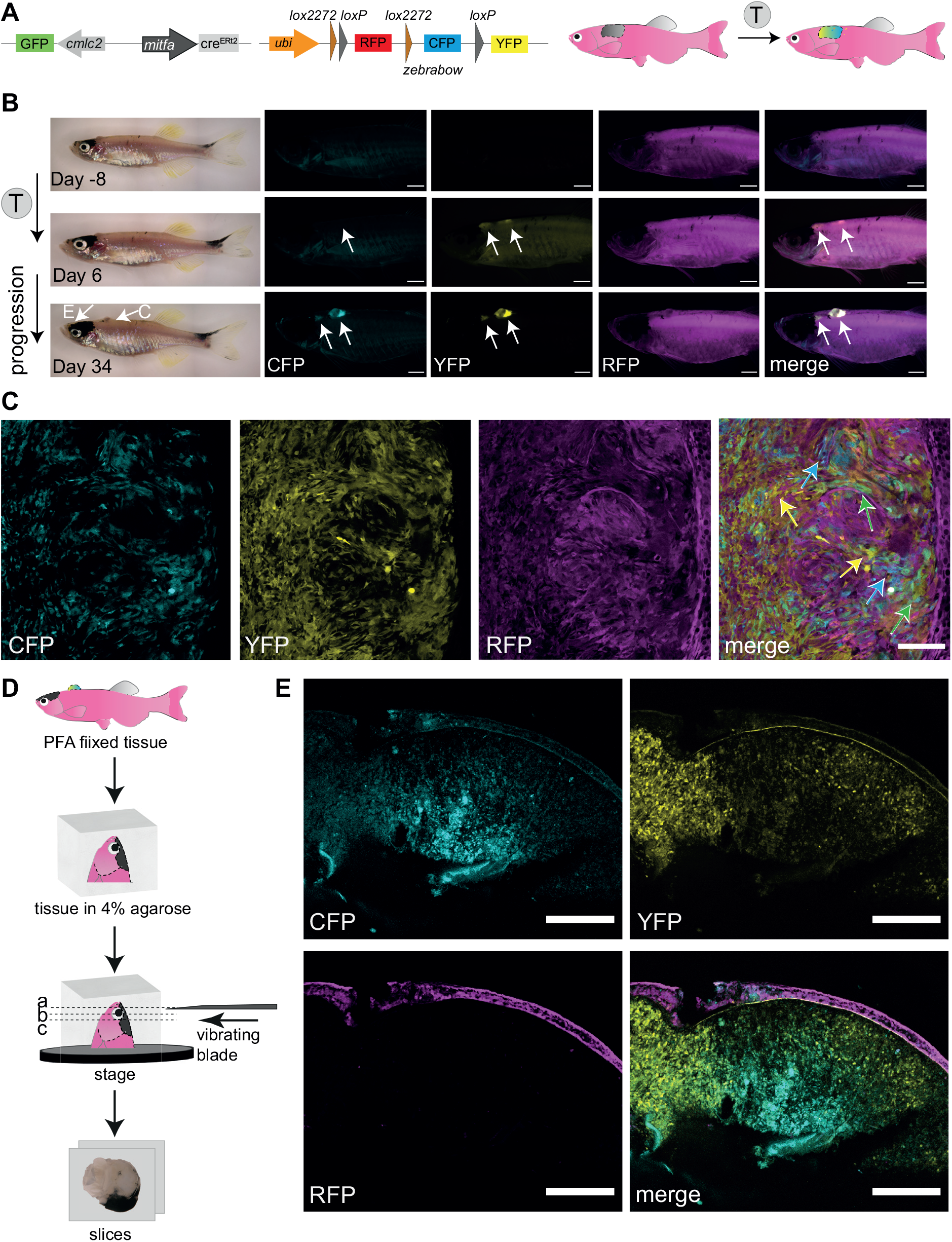
Successful fluorophore switch using the zebrabow system in adult zebrafish primary melanoma following tamoxifen treatment. **A.** A schematic of *mitfa:cre^ERt2^* and *ubi:zebrabow* constructs (Pan *et al.,* 2013) which enables recombination from red signal (RFP) to a stochastic combination of cyan (CFP), yellow (YFP) and red (RFP) signal in *mitfa:cre^ERt2^* expressing cells after tamoxifen (T) treatment. The *cardiac myosin light chain (cmlc) 2* promoter drives *GFP* expression and facilitates screening for the *cre^ERt2^* line based on GFP+ heart myocardium. **B.** Multicolour labelling in adult zebrafish melanomas. An adult zebrafish with two primary melanomas 8 days before the tamoxifen (T) treatment course (top), and 6 days and 34 days after the start of the treatment. Increasing *de novo* expression of CFP and YFP proteins in both pigmented and unpigmented tumours can be detected after tamoxifen treatments (4 μM). N = 3 fish, Scale bar = 1 mm, white arrows point to two tumour locations. C and E labels tissue images in panels C and E respectively. **C.** Confocal multicolour imaging of zebrafish melanoma. Single z-plane of confocal acquisition of tamoxifen (T) treated fish shows individual cells acquiring varied combinations of CFP, YFP and RFP (blue, green and yellow arrows on merged image). N = 3 fish, scale bar = 100 μm. **D.** Overview of the vibratome sectioning protocol. The PFA fixed tissue is mounted in 4% agarose and cut using vibrating blade into 400 μm thick sections to capture the pigmented tumour (E). **E.** Vibratome section imaging of a tamoxifen-treated fish (tissue location as shown in **B**). Clusters of CFP, YFP or double expressing cells are clearly visible in areas of the pigmented tumour. STD projections of confocal z-stacks of a PFA fixed sectioned tissue. N = 3 fish, scale bar = 200 μm. See also **Figure S1B-D**.

Melanoma tissue often varies in the pigmentation level between individual tumours, and while non-pigmented tumours permit direct detection by fluorescence, intense pigmentation can obscure the fluorescent signal. To overcome this challenge, we performed vibratome sectioning of PFA fixed melanoma tissue (**Fig. 3D**) followed by confocal microscopy. These imaging data show that combining thick tissue sectioning with confocal microscopy permits detection of CFP and YFP channels even in highly pigmented tissues (**Fig. 3E, Fig. S1D**). While we did not evaluate clonal evolution in melanoma progression (as it would require more rigorous analysis of colour switch stochasticity), this experiment demonstrated the potential for our tamoxifen treatment protocol for application in clonal analysis studies in zebrafish cancer models.

### Melanoma persister cells originate from the primary tumour

Next, we used our conditional MITF-dependent BRAF^V600E^ p53^M214K^ *ubi:switch* model to trace melanoma cells from the primary tumour as it regresses and thereby determine if the cells detected at the MRD site originate from the primary tumour (**Fig. 4A**). Tamoxifen treatment of early-stage melanoma resulted in a GFP-to-mCherry switch in tumour lesions that was absent in DMSO-treated animals (**Fig. 4B-C**). Following a period of tumour growth, we transferred fish to a higher temperature to turn off MITF activity and cause tumour regression until no melanoma was detectable morphologically (5-10 weeks; see Methods). Strikingly, at the MRD site, we could detect mCherry+ persister cells in tamoxifen-treated fish (**Fig. 4B-C**). Confocal microscopy and quantification confirmed that tamoxifen-treated fish showed significantly greater mCherry signal at the MRD site compared to DMSO-treated fish (**Fig. 4D-E**). Whilst most DMSO-treated fish did not show any mCherry signal, we could detect it in a small proportion of the control fish (**Fig. 4E**). This may be due to *cre* expression (leakage) in those tissues where *mitfa* expression is very high (for example in some nodular tumours as we have described previously (Travnickova *et al.,* 2019)). Immunostaining of regressed melanoma tissue sections confirmed the presence of mCherry+ persister cells at the MRD site (**Fig. 4F**). These data indicate that primary melanoma cells directly give rise to persister cells at the MRD site.

**Figure 4:**
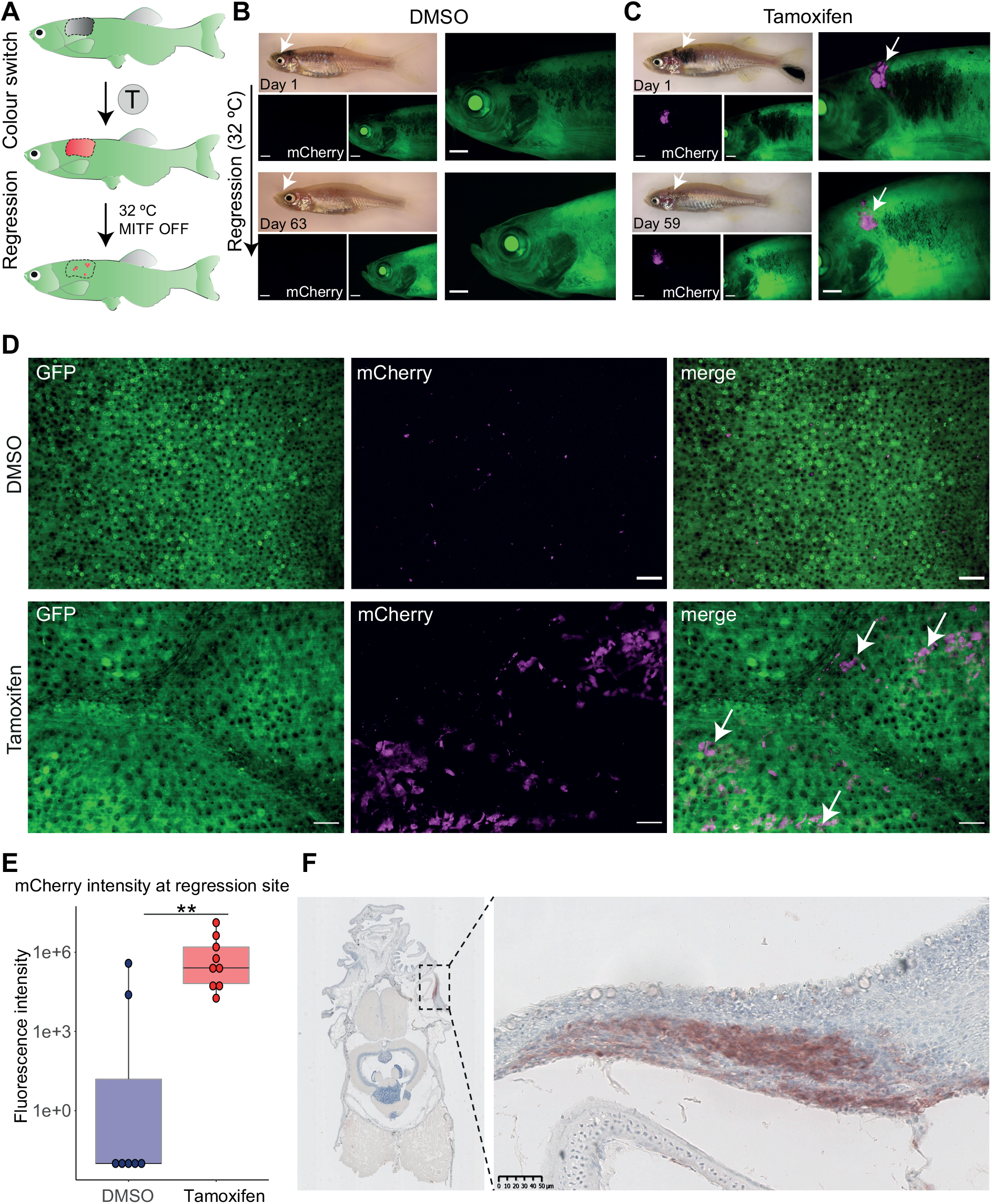
Persister cells originate from the primary tumour. **A**. The experimental workflow of tamoxifen (T) treatment and melanoma regression at the higher water temperature (32°C, MITF OFF). At the higher temperature *mitfa* RNA is still expressed but is not spliced correctly and therefore MITF activity is abolished. Loss of MITF activity leads to tumour regression with remaining persister cells at the minimal residual disease (MRD) site. **B**. A fish treated with vehicle (DMSO 0.05%) showing no green-to-red recombination in the primary tumour (top) or after regression (bottom). Scale bar = 1 mm, white arrow points at the tumour site, N = 4 fish. **C**. A fish treated with tamoxifen (5 μM) showing expression of mCherry positive cells in a primary tumour (magenta, white arrows) that are still detected after 59 days after initiation of melanoma regression (bottom, white arrows). Scale bar = 1 mm, N = 7 fish. **D**. STD projections of confocal z-stack acquisitions of regression sites of fish treated with DMSO or Tamoxifen. White arrows point to mCherry positive cells (magenta) only present in the tamoxifen treated condition. Scale bar = 50 um, DMSO: N = 4 fish; Tamoxifen: N = 7 fish. **E.** Box plot of fluorescence intensity quantification of mCherry signal using average intensity projections of confocal images of residual disease. DMSO: N = 4 fish with 7 regression sites; Tamoxifen: N = 7 fish with 9 regression sites; ** p<0.01, Wilcoxon test. **F.** Immunohistochemistry of the MRD site. FFPE transverse section stained with anti-mCherry antibody reveals “switched” cells at the MRD site. Right panel shows an enlarged view of MRD, outlined by the black dashed rectangle (left panel). Signal in red (AEC substrate) counterstained with nuclear marker haematoxylin (blue). Scale bar= 50 μm, N = 3 fish.

### Melanoma persister cells directly contribute to tumour recurrence

Next, we asked if the tamoxifen-induced GFP-to-mCherry switch could also be applied to melanoma persister cells in the MRD site (**Fig. 5A**). We transferred our melanoma fish to a higher water temperature to prevent the correct splicing of *mitfa* (and to thereby turn off Mitfa protein activity) to cause melanoma regression. Because the *mitfa^vc7^* mutation is an RNA splicing mutation, the expression of *mitfa* or reporters under the control of *mitfa* (e.g. *mitfa:GFP; mitfa:cre^ERt2^)* are not affected at the restrictive temperature (**Fig. 5B**). Once the melanoma had fully regressed, we treated the fish with tamoxifen. We detected mCherry+ cells 6 days after the start of tamoxifen treatment at the regression site (**Fig. 5C-D**). Using confocal microscopy, we were able to validate the presence of mCherry+ cells in the tamoxifen-treated group (**Fig. 5E**), that were morphologically similar to our previous observations using *Tg(mitfa:GFP)* transgenic line (Travnickova *et al.,* 2019).

**Figure 5:**
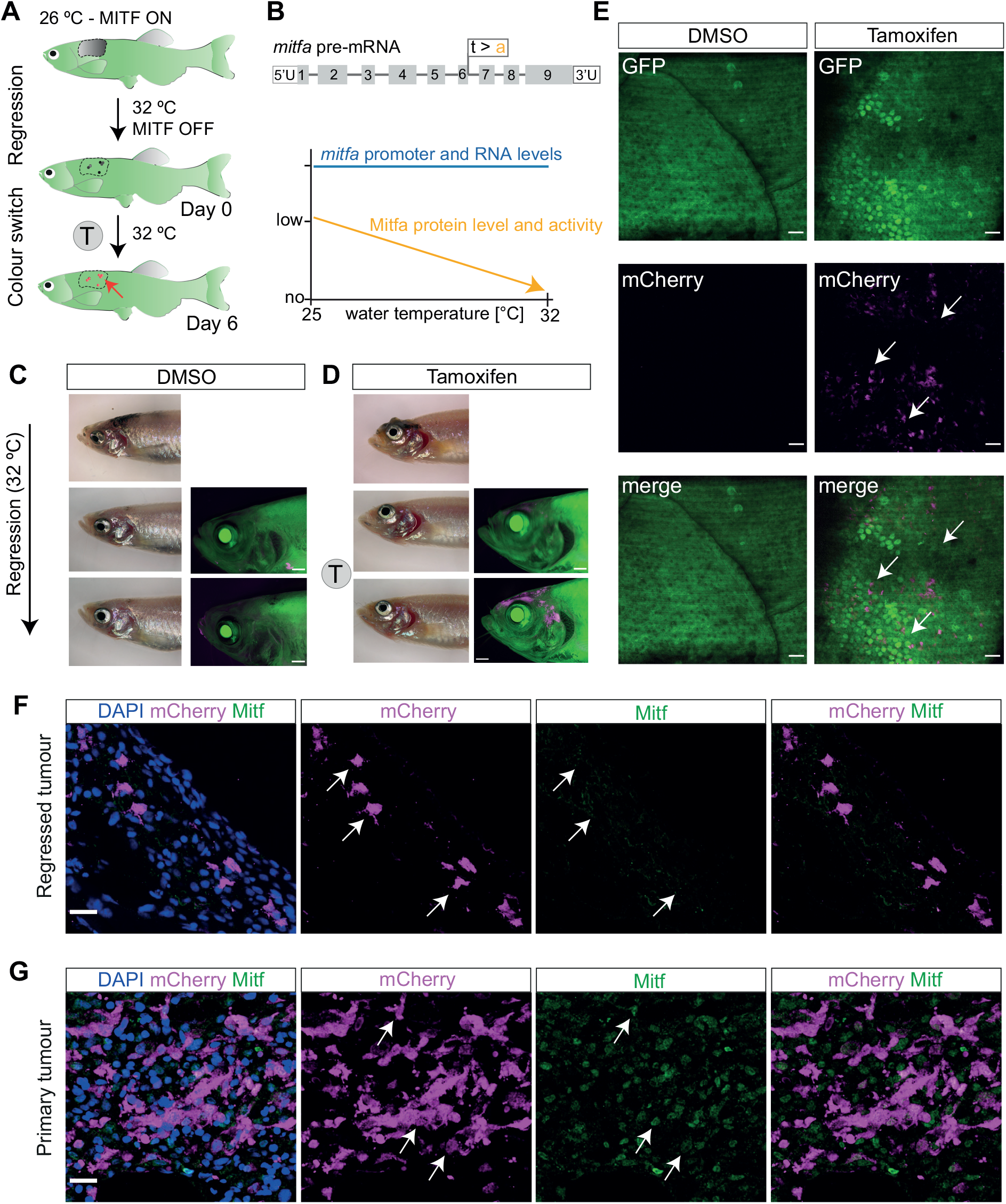
Successful GFP to mCherry fluorophore switch in persister cells. **A.** Schematic representation of the experimental workflow of tamoxifen (T) treatment following melanoma regression. The red arrow points to the location of recombined melanoma persister cells at MRD site. **B.** Overview of *mitfa^vc7^* mutant controlling Mitfa activity. (Top) Schematic of *mitfa* pre-RNA with the location of *vc7* mutation at the end of exon 6. (Bottom) Graphical overview of effect of *mitfa^vc7^* mutation on Mitfa protein level and activity (orange) at increasing water temperatures. Mitfa protein and activity levels decrease with increasing water temperature, unlike the *mitfa* RNA levels which remain expressed (blue). Schematics adapted from (Travnickova *et al.,* 2019). **C.** A representative image of a control fish with no mCherry positive cells detectable at the MRD site without tamoxifen treatment. Following melanoma regression and subsequent treatment with DMSO (0.04% at 31-32°C) there is no green-to-red recombination in the melanoma MRD (bottom panel) compared to pre-treatment (middle panel). Scale bar = 1 mm, N = 3 fish. **D.** A representative image of a tamoxifen-treated fish with newly ‘switched’ mCherry positive cells (magenta) detectable at the MRD site. Following melanoma regression and subsequent treatment with tamoxifen (4 μM, at 31-32°C) there is specific green-to-red recombination in the melanoma MRD (bottom panel) compared to pre-treatment (middle panel). Scale bar = 1 mm, N = 3 fish. **E.** STD projections of confocal z-stack acquisitions showing MRD sites in fish treated with DMSO or Tamoxifen. White arrows indicate clusters of mCherry positive cells present only in tamoxifen treated condition. Scale bar = 50 μm, N = 3 fish. **F.** Melanoma cells at the MRD site express mCherry, but lack Mitfa protein (white arrows). STD projections of confocal z-stack acquisitions of immunofluorescence staining of the tamoxifen-treated MRD with antibodies for mCherry and Mitfa proteins and with DAPI nuclear staining. Scale bar = 15 μm. **G.** Melanoma cells in the primary tumour express both mCherry and Mitfa protein (white arrows). STD projections of confocal z-stack acquisitions of immunofluorescence staining of the tamoxifen-treated primary tumour, showing staining of mCherry and Mitfa proteins with DAPI nuclear staining Scale bar = 15 μm. See also **Figure S2.**

We have previously demonstrated that melanoma persister cells in the MRD site do not express Mitfa protein (called MITF-independent), but maintain a neural crest identity and express Sox10 (Travnickova *et al.,* 2019). To evaluate if the ‘switched’ mCherry positive cells retain the same properties, we immunostained the primary and regressed tamoxifen-treated melanoma tissue sections with mCherry and Mitfa antibodies (**Fig. 5F-G**). No Mitfa expression was detected in ‘switched’ mCherry positive melanoma cells in the regressed tumour. In contrast, the primary tumour showed abundant Mitfa expression in mCherry ‘switched’ cells (**Fig. 5F-G**). Consecutive sections of the MRD site were stained with mCherry and Sox10 antibodies and confirmed that the mCherry positive cells were Sox10+ persister melanoma cells (**Fig. S2**). These data show that ‘switched’ mCherry positive persister cells at the MRD site are MITF-independent.

Melanoma persister cells have been proposed to contribute to disease recurrence and drug resistance (Marin-Bejar *et al.,* 2021; Rambow *et al.,* 2018b; Shen et al., 2020a; Travnickova *et al.,* 2019; Vendramin et al., 2021), but this has not been demonstrated in an animal model using lineage tracing. Having validated the successful GFP-to-mCherry switch at the MRD site, we next sought to follow the persister cells during recurrence. Once again, we performed tamoxifen treatment and detected GFP-to-mCherry switched persister cells at the MRD site. We then restored MITF activity by lowering the water temperature and followed melanoma recurrence. Over 50 days, the mCherry+ cells continuously increased concomitantly with melanoma recurrence (**Fig. 6A-B)**. Fluorescence area quantification of individual fish MRD sites show that mCherry+ cells drive tumour growth over time (**Fig. 6C**). This demonstrates that persister cells directly contribute to the growth of recurrent disease.

**Figure 6:**
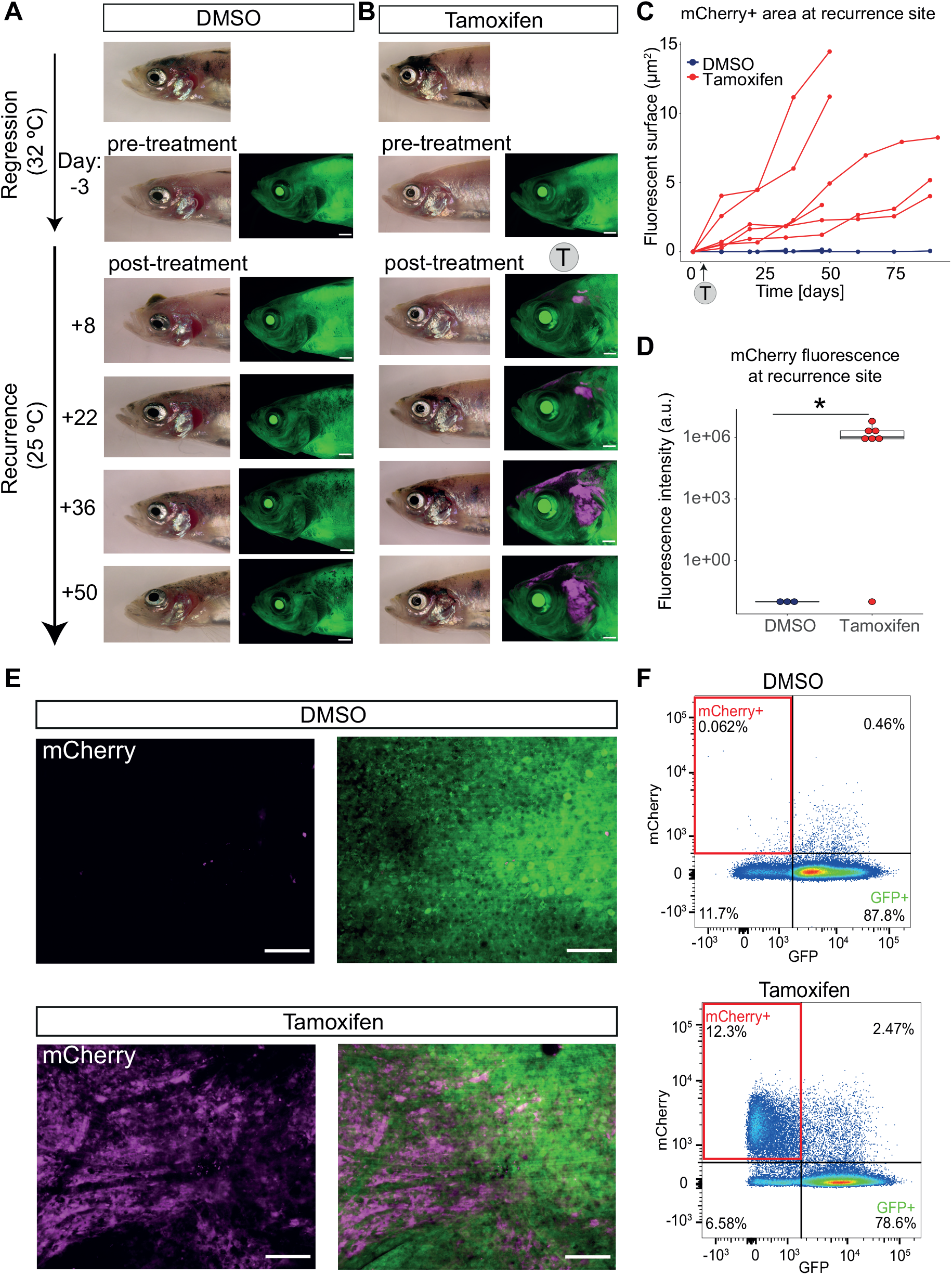
Persister cells directly contribute to melanoma recurrent disease. **A.** Representative images of control fish with no mCherry fluorescence. A fish with a regressed melanoma (top image: primary tumour; below: regressed tumour) treated with vehicle (DMSO 0.04-0.05% at 32°C) showing no green-to-red fluorophore switch in the melanoma MRD compared to the pre-treatment image. Follow–up bright field and fluorescent acquisitions (every 14 days) show progression of tumour recurrence over time with no appearance or increase of mCherry fluorescence. Scale bar = 1 mm. N= 5 fish (sum of two independent experiments). **B.** mCherry positive persister cells from MRD are present through to recurrence. A fish with a regressed melanoma (top image: primary tumour; below: regressed tumour) treated with 4-5 μM tamoxifen (at 32°C) showing green-to-red fluorophore switch (in magenta) in the melanoma MRD. Follow–up brightfield and fluorescent acquisitions (every 14 days) show progression of tumour recurrence over time with increasing mCherry signal. Scale bar = 1 mm. N = 7 fish (sum of two independent experiments). **C.** mCherry positive persister cells increase over time. Line plot of area quantification of mCherry signal using fluorescent acquisitions over time during melanoma recurrence. DMSO (blue): N = 5 fish; Tamoxifen treated group (red): N = 6 fish (sum of two independent experiments for both groups). Trajectories of melanoma growth represent individual fish. **D.** Box plot of fluorescence intensity quantification of mCherry signal of recurrent disease that has grown from MRD treated with tamoxifen. Intensity was measured using average intensity projections of confocal images of recurrence sites comparing DMSO to Tamoxifen condition. DMSO: N=3 fish; Tamoxifen treated group: N=4 fish with 7 recurred melanomas * p<0.05, Wilcoxon test. Outlier sample in Tamoxifen treatment group is likely to be false negative as mCherry fluorescence was later confirmed using flow cytometry in this highly pigmented sample. **E.** mCherry is present only in recurrent disease that has grown from MRD treated with tamoxifen. STD projections of z-stack confocal acquisition show recurred melanoma sites of fish treated with vehicle (DMSO, top row) or Tamoxifen (bottom row) at the MRD stage. Scale bar = 100 μm. **F.** mCherry+ cells in recurrent disease are detectable by FACS. A representative pseudocolour flow cytometry plot of a recurred melanoma lesion from DMSO treated fish and tamoxifen treated fish. mCherry positive cells highlighted in red rectangle (12.3% vs 0.062%). DMSO: N= 4 fish; Tamoxifen: N = 5 fish (sum of two independent experiments).

Confocal microscopy enabled us to visualise the tumours at cellular resolution, showing fields of mCherry+ melanoma cells in recurrent disease (**Fig. 6D-E**). Quantification of the mCherry+ fluorescence indicated that most recurrent disease sites expressed mCherry (**Fig. 6D**; 6/7 recurrence sites with an average intensity of 2×10^5^ a.u.). For one sample, we found the fluorescence was hindered by strong pigmentation. Given this, we validated the presence of mCherry+ cells in this and other recurrent tumours using flow cytometry and could clearly detect mCherry even in highly pigmented samples (**Fig. 6F**).

### Transcriptional plasticity of melanoma persister cells in recurrent disease

We hypothesised that persister cells directly lead to recurrent disease by transitioning from an MITF-independent to a Mitfa+ state. As previously described and shown here (**Fig. 5F-G**) persister cells do not express Mitfa protein, but maintain their expression of Sox10 (Travnickova *et al.,* 2019). We applied multiplex immunohistochemistry (MIHC) on sections of the recurred tumours to determine the protein expression levels of Mitfa and other melanoma markers within recurrent disease (**Fig. 7A-D**). MIHC allows sequential staining and stripping of several antibodies on the same slide (Pirici et al., 2009). We used an antibody against the mCherry protein to label the “switched” cells in tamoxifen treated fish compared to DMSO control, together with antibodies against the melanoma markers Sox10 and Mitfa (**Fig. 7B-D**). As anticipated, recurred tumours from DMSO controls lacked mCherry signal in Sox10+ and Mitfa+ tumour cells (**Fig. 7B, B’**). By overlaying the images of mCherry staining with Sox10 and Mitfa staining we could demonstrate that the mCherry+ cells within the tamoxifen treated recurred tumour expressed both Sox10 and Mitfa (**Fig. 7D’, Fig. S3**). These results indicate that melanoma persister cells undergo a cell state switch from MITF-independent to Mitfa+ expressing cells, regaining features of the primary tumour, and providing direct evidence of tumour cell transcriptional plasticity *in vivo.*

**Figure 7:**
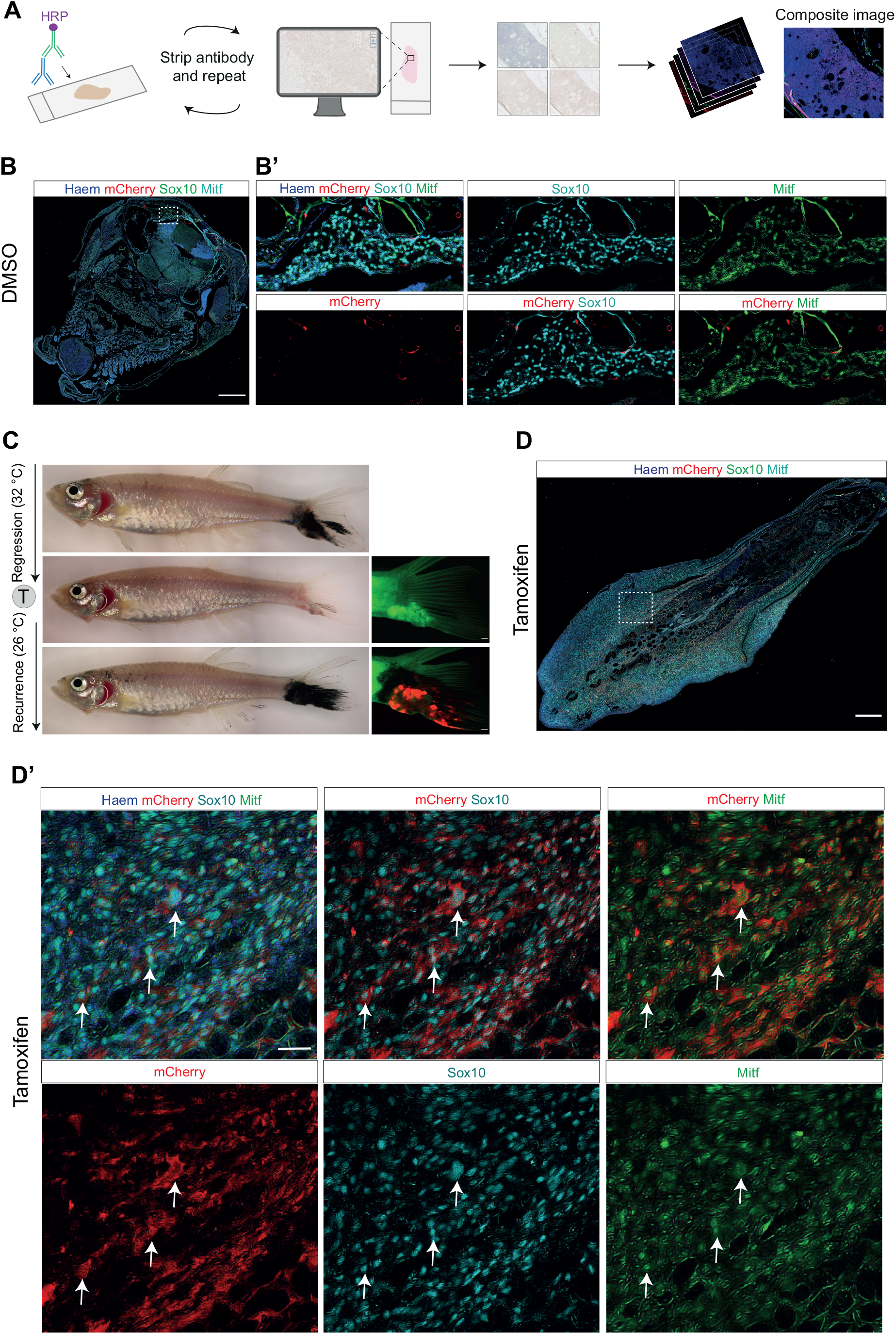
MITF-independent persister cells regain Mitfa protein expression in recurrent disease. **A.** Overview of the multiplex immunohistochemistry protocol. The antibody staining is repeatedly imaged and stripped, images are pseudocoloured and stacked on ImageJ to generate a composite image. **B.** DMSO-treated melanoma recurrent disease on top of the head, labelled with Sox10, mCherry and Mitfa antibodies overlaid with haematoxylin counterstain; dashed white box indicates area of section shown in **B’**. Scale bar, 250 μm. **B’.** Enlarged views of DMSO-treated tumour section shown in **B**, showing lack of mCherry staining in melanoma cells positive for Sox10 and Mitfa. Scale bar, 50 μm. Observable mCherry signal represents trapped antibody in loose tissue fragments. **C.** (Left, top to bottom) brightfield images of melanoma tail fin tumour through growth, regression and recurrence. (Right, middle and bottom) fluorescent images of regressed and recurred melanoma tail fin tumour, showing mCherry fluorescence in the recurred tumour following tamoxifen mediated recombination in the MRD. Scale bars, 500 μm. **D**. Recurrent disease from MITF-independent melanoma persister cells regain Mitfa protein expression. Overview of tamoxifen treated melanoma recurrent disease in the tail fin, showing Sox10, mCherry and Mitfa protein staining overlaid with haematoxylin counterstain, dashed white box indicates area of section shown in **D’**. Scale bar, 250 μm. **D’.** Enlarged views of tumour section shown in **D**, showing individual staining of Sox10, Mitfa, and mCherry proteins. Scale bar, 25 μm. White arrows highlight cells positive for Sox10, Mitfa and mCherry staining. See also **Figure S3**.

## DISCUSSION

Transcriptional heterogeneity underlies the high levels of tumour cell plasticity in melanoma. Here, we apply fate mapping and MIHC to tumour cell populations through melanoma disease states to show that (1) persister cells originate from the primary tumour; (2) persister cells directly contribute to melanoma recurrence; and (3) persister cells exhibit plasticity from a MITF-independent state to Mitfa expressing melanoma cells in recurrent disease. Thus, persister cells will be a critical drug target for delaying (or even preventing) melanoma recurrent disease.

Our system offers the opportunity to understand how individual cell states, and indeed even individual cells, contribute to drug resistance and tumour progression in zebrafish cancer models. Lessons from mouse models indicate that the efficiency of recombination is largely dependent on the expression level and pattern of the chosen promoter and thus the tamoxifen protocol often needs to be adapted for each *cre^ERt2^* line (Ellisor and Zervas, 2010; Jahn et al., 2018). In particular, the Bally-Cuif group recently established a 4-OHT protocol for mosaic recombination using short-term drug exposure and long-term multiple-day immersion for maximal recombination, similar to our approach (Than-Trong *et al.,* 2020). Hence, as inducible lineage tracing methods become more widely applied in zebrafish adults, the zebrafish community will benefit from ensuring tamoxifen protocols for each promoter are easily accessible.

Here, we showed the fate mapping of the *mitfa:cre^ERt2^* expressing melanoma cell population through disease stages, which represents a major portion of melanoma cells in our model. MITF is one of the universal diagnostic markers of cutaneous melanoma based on its expression in most melanoma cells (Compton et al., 2015). Previously, we and others have demonstrated the level and importance of transcriptional heterogeneity not only in the primary tumour but also in the persister cells at the MRD site (Marin-Bejar *et al.,* 2021; Rambow *et al.,* 2018b; Shen *et al.,* 2020b; Travnickova *et al.,* 2019; Wouters et al., 2020). The persister cell states in our zebrafish models share conserved mechanisms with human melanoma cells depleted for MITF or following BRAF inhibitor drug treatment (Dilshat et al., 2021; Rambow et al., 2018a; Travnickova *et al.,* 2019). The lineage tracing system we present here has the potential to enable fate mapping of different transcriptional cell states at the MRD site and evaluate their contribution to disease progression and response to drug treatment (Lu and Patton, 2022). This can be achieved using cell state specific markers to recombine a subpopulation of the tumour at the desired stage.

In conclusion, we have successfully applied and validated an inducible *cre^ERt2^/loxP* lineage tracing system in an adult zebrafish melanoma model to follow cell states through disease stages. Combining fate mapping with live imaging, IHC and MIHC, we provide direct evidence for the contribution of cells at the MRD site to tumour recurrence, whilst simultaneously demonstrating their MITF-independent to Mitfa+ cell state shift. This system has the potential for widespread usage in the study of cancer cell plasticity over time and in the evolution of therapy resistance.

## METHODS

### Experimental Models and Husbandry

Zebrafish were maintained in accordance with UK Home Office regulations, UK Animals (Scientific Procedures) Act 1986, amended in 2013, and European Directive 2010/63/EU under project license 70/8000 and P8F7F7E52. All experiments were approved by the Home office and AWERB (University of Edinburgh Ethics Committee).

Fish stocks used were: *mitfa^vc7^* (Johnson *et al.,* 2011; Zeng *et al.,* 2015), *Tg(mitfa-BRAF^V600E^), tp53^M214K^(lf)* (Patton et al., 2005)*, Tg(ubb:loxP-EGFP-loxP-mCherry)* (referred as *ubi:Switch)* (Mosimann *et al.,* 2011), *Tg(ubb:lox2272-loxP-Tomato-lox2272-Cerulean-loxP-YFP)* (referred here as *Tg(ubb:lox2272-loxP-RFP-lox2270-CFP-loxP-YFP)* or *ubi:zebrabow);* (Pan *et al.,* 2013), *Tg(mitfa:cre^ERt2^)* (this study). Combined transgenic and mutant lines were generated by crossing. Adult fish were maintained either at 28.5°C, 25°C or 32°C under 14:10 light:dark cycles.

### Generation of zebrafish transgenic line *mitfa:cre^ERt2^*

Cre^ERt2^ was amplified by PCR using *pCGA_cre^ERt2^* (Addgene plasmid #14797) as template. A nuclear localization signal (NLS) was added to its N terminal. The gateway primer sequences are *cre^ERt2^gateF* 5’-GGGGACAAGTTTGTACAAAAAAGCAGGCTGCCACCATGCCCAAGAAGAAGAGGAAGGT GTCCAATTTACTGACCGTACACC-3’ and *cre^ERt2^gateR:* 5’GGGGACCACTTTGTACAAGAAAGCTGGGTTCAAGCTGTGGCAGGGAAAC-3’. The PCR product was cloned into *pDONOR221* resulting in *pME-cre^ERt2^,* a middle entry vector. cre^ERt2^ was then cloned together with the 2.1 kb zebrafish *mitfa* promoter into *pDestTol2CG2* destination vector by the Tol2kit gateway cloning method (Kwan et al., 2007), resulting in the *pEXPmitfa-cre^ERt2^* expression vector. For selection purposes, the construct contains an additional GFP coding sequence expressed from the heart specific *cmlc2* promoter. Two nl of mixed *pEXPmitfa-cre^ERt2^* plasmid and *Tol2* mRNA (25 ng μl^-1^ and 35 ng μl^-1^ respectively) were injected into 1-cell stage zebrafish embryos. Zebrafish embryos with the GFP transgenic marker in the heart were selected and grown to adulthood, then bred with wildtype fish to establish stable lines

### Genotyping

Zebrafish were genotyped using DNA extracted from fin clipped tissue using DirectPCR lysis reagent (Viagen) complemented with 0.1 mg ml^-1^ proteinase K. Polymerase chain reaction (PCR) was used to establish the mutant allele status *p53^M214k^* and to verify the presence of transgene *mitfa-BRAF^V600E^* as described in detail before (Wojciechowska et al., 2016).

### Temperature-controlled melanoma regression and recurrence

To induce melanoma regression, selected adult melanoma-prone fish homozygous for *mitfa^vc7^* mutation were transferred to tanks with heaters that kept the water temperature at 32°C compared to standard system temperature of 28.5°C. Fish were monitored twice a week for tumour changes and imaged every two weeks to visually compare the regression over time. A tumour was considered fully regressed once no melanoma was visually detectable and stayed unchanged during two consecutive time points of monitoring (usual range of regression being 5-10 weeks). To allow melanoma recurrence, regressed fish were transferred to tanks at room temperature (24-26°C), while monitored daily for tumour progression and health condition of the fish. Fish were always imaged prior to transfer from one temperature to another, then imaged regularly to track their progression and monitored daily. Each fish was followed through recurrence until the time point the tumour reached a similar size to the primary tumour or until reaching the humane end point criteria based on general fish fitness, swimming behaviour and location of tumour (with a maximum of 3 months of recurrent disease growth).

The emergence of melanoma ranges between 2 and 14 months of age. All fish admitted to the experiment were over 3 months old and all experiments were finished before fish reached 18 months of age. Both females and males were included in the experiments: on average, across experiments, the ratio was 58% females (50-66%) and 42% males (33-50%).

### Tamoxifen and 4-hydroxytamoxifen treatment

The tamoxifen treatment protocol was adapted from Gemberling *et al* and Pinzon-Olejua *et al.* (Gemberling et al., 2015; Pinzon-Olejua et al., 2017). The tamoxifen stock solution (Cayman Chemicals or Fisher Scientific) was prepared by dissolving tamoxifen powder in DMSO to reach 10mM concentration. We prepared fresh stock for each round of treatment on day 0 for the whole 3-day-treatment course and stored at −20°C in 200-250 μl aliquots. Tamoxifen stock solution (or corresponding DMSO volume as a control) was diluted with 500 ml E3 medium (5mM NaCl, 0.17 mM KCl, 0.33 mM CaCl_2_, 0.33 mM MgSO_4_) in a fish carrier tank and fish immersed in the solution for 11-12 hours overnight (protected from light using a black chamber box or by wrapping the tank in aluminium foil) for 3 consecutive nights, with a drug pause during the day time. For the tamoxifen treatment of fish with regressed melanoma, the carrier tanks with fish immersed in the tamoxifen or DMSO were kept in a water bath set at 31°C to ensure the continuous suppression of MITF activity.

4-hydroxytamoxifen (4-OHT, Sigma H6278) stock was prepared by dissolving the powder in ethanol to reach 5 or 10 mM concentration and thoroughly vortexed and aliquoted into 100 μl aliquots. The 4-OHT stock solution was then diluted into 200 μM intermediate stock in E3, vortexed thoroughly while protected from light and then diluted in E3 to the final working concentration 20 μM. Embryos were placed into 6-well-plates with 3 ml 20 μM 4-OHT, protected from the light and incubated at 28°C. The 4-OHT was refreshed daily for a total incubation of 3 days. Embryos were then washed three times with E3 and incubated in E3 prior to imaging. In **Figures 1D-E**, embryos were incubated for 6 hours after the end of 4-OHT treatment.

### Live imaging of adult zebrafish

Fish were briefly anaesthetised in tricaine solution (MS-222, 0.1 g l^-1^), placed onto a petri dish while immersed in the tricaine solution and imaged using a Nikon COOLPIX5400 camera attached to a brightfield microscope (Nikon SMZ1500). Each fish was imaged using 3 consecutive photo shots that were later merged together using Adobe Photoshop automerge function. For fluorescent imaging, anaesthetized fish were imaged under the Leica MZFIII stereomicroscope equipped with Retiga R1 camera operated via μManager software (Edelstein et al., 2010). Images were taken using GFP and mCherry fluorescent filters. The fluorescent images were processed using Fiji software version 2.1.0 (Schindelin et al., 2012) for pseudocolouring and adjusting minimal and maximal intensity.

For zebrabow live imaging, anaesthetised fish were imaged under the Leica M205FCA stereoscope equipped with Leica K8 camera and operated via LASX software (Leica). Images were taken with 0.63x objective at 1x or 2.5x zoom using CFP, YFP and mCherry fluorescent filters and processed as above. As shown in **Figure S1B,** the CFP channel showed a certain level of autofluorescence, most prominent in the creases of jaw and gills that was not considered as positive signal.

### Zebrafish tissue and embryo confocal imaging

For confocal microscopy, zebrafish embryos were anaesthetised with MS-222 0.1 g.l^-1^ and adult fish were culled by overdose of anaesthetic (MS-222 1 g l^-1^, followed by death confirmation) and embedded in 1% low melting point (LMP) agarose (Invitrogen) with the tumour or regressed tissue oriented to the bottom of 6-well glass bottom plate (Cellvis). For dorsal imaging of zebrafish embryos, fish were incubated for 5 min with 5 mg.ml^-1^ adrenaline (epinephrine, Sigma, E4642) in order to contract the pigment in melanocytes prior mounting in LMP for imaging. Images of *ubi:Switch* transgenic were obtained under Andor Dragonfly spinning disk confocal microscope equipped with Andor Zyla sCMOS camera through 20x air objective with 2048×2048 px resolution and 1 μm interval. Images of *ubi:zebrabow* transgenic were obtained under Leica STELLARIS 8 confocal microscope equipped with white light laser through 10x and 20x air objectives with 1024×1024 px resolution and 1μm (20x) or 3 μm (10x) intervals. In contrast to stereoscope imaging, we did not detect CFP channel autofluorescence using confocal microscopy.

### Image processing and Fluorescence quantifications

Confocal acquisitions were processed using Fiji software version 2.1.0 (Schindelin *et al.,* 2012). Standard deviation intensity projections of all z-slices were used except for fluorescence intensity quantification where average intensity projections were used. Fluorescence intensity has been measured as described (McCloy et al., 2014). Briefly, average intensity projections of z-stack acquired using Andor Dragonfly spinning disk confocal microscope were split by channel to select mCherry only. Intensity was then measured of the whole imaged region and small region without any tissue used to determine background. To calculate fluorescence intensity, corrected total cell fluorescence (CTCF) was calculated by subtracting the background fluorescence intensity. Any negative value was replaced by 0. Fluorescence area over time was measured on the mCherry channel of Leica MZFIII epifluorescence microscope acquisitions taken before and after the tamoxifen treatment and then every 2 weeks through melanoma recurrence. Area was manually depicted using the polygon selection tool in Fiji and then measure in μm^2^. Any sample that showed fluorescence signal prior the tamoxifen treatment was excluded from the analysis. Graphs summarizing the fluorescence intensity quantification were made using R software version 4.0.3 via R studio interface version 1.1.456 equipped with ggplot2 package (R-Core-Team, 2020; RStudio-Team, 2016; Wickham, 2016).

### Vibratome sectioning

Fish were fixed overnight in 4% PFA at 4°C with agitation. The piece of fish tissue was then rinsed twice with PBS for several minutes and mounted in 4% agarose/PBS using a well from a 6-well-plate as a mold. Any excess agarose above 0.5 cm from the tissue was discarded prior sectioning. Tissue was sectioned transversely using Leica VT1200S vibratome at a thickness of 400 μm and the speed of vibrating blade was 0.1 mm s^-1^. Sections were collected in PBS and mounted in 1% LMP agarose in a 6-well glass bottom plate prior imaging at Leica STELLARIS 8 as described above.

### Fluorescence activated cell sorting

Adult tamoxifen treated or DMSO control fish with recurred tumours were culled using overdose of tricaine (MS-222, 1 g l^-1^ followed by death confirmation). Tumours were individually dissected and chopped using sterile No.9 blade scalpels. Samples were then dissociated with 0.25 mg ml^-1^ liberase TL (Roche) for 15 minutes at room temperature while inverting the tube, re-suspended in FACSmax cell dissociation solution (Genlantis) and filtered through 40μm cell strainer. Samples were sorted by a FACSAria2 SORP instrument (BD Biosciences UK) equipped with 405nm, 488nm and 561 nm lasers. Green fluorescence was detected using GFP filters 525/50 BP and 488 nm laser, red fluorescence using mCherry filters 610/20 BP and 561 nm laser and live cells selected as DAPI negative using DAPI filters 450/20 BP and 405 nm laser. Data were analyzed using FlowJo software (BD Biosciences) version 10.8.1.

### Histology

Fish samples were collected, fixed and processed as described in detail in (Wojciechowska *et al.,* 2016). Briefly, tissue was fixed by immersion in 4% PFA at 4°C with agitation for 3 days, decalcified in 0.5 M EDTA (pH 8) at 4°C with agitation for 5 days and then stored in 70% ethanol at 4°C. To obtain sections for pathology analysis, tissue was processed in 95% ethanol, absolute alcohol, xylene and paraffin, embedded in blocks, cut into 5 μm thick sections and placed onto glass slides. Haematoxylin and eosin staining and immunohistochemistry were performed as described in detail in (Wojciechowska *et al.,* 2016). The slides were imaged using Hamamatsu Nanozoomer SlideScanner and the images were processed using NDP.3 software.

### Immunofluorescence staining

FFPE sections were deparaffinised in xylene and gradually rehydrated through baths of ethanol of decreasing concentrations. Sections were bleached in 1% KOH/3% H_2_O_2_ solution for 15 minutes prior to subjected to heat mediated antigen retrieval in citrate buffer (0.01 M, pH6). Sections were blocked in 10% goat serum for 1 hour and incubated with following primary antibodies: mCherry (Abcam, ab125096, dilution 1:3,000 or 1:4,000) (Kobayashi et al., 2014), Mitfa (Sigma, SAB2702433, 1:1,000) and Sox10 (Abcam, ab125096, 1:4,000) (Travnickova *et al.,* 2019) overnight at 4 °C. After PBS/0.1% tween washes, sections were incubated with 488 (# A11001) or 546 (# A11003) AlexaFluor Goat anti-mouse and/or 647 (#A31573) AlexaFluor Goat anti-rabbit secondary antibodies at 1:500 dilution (Invitrogen) and DAPI (Sigma, 1:500 dilution) for 30 min in dark at room temperature and mounted in hydrophilic mounting medium prior imaging. Sections were imaged using Andor Dragonfly confocal microscope equipped with Andor Zyla sCMOS camera using 20x and 40x air objectives.

### Multiplex Immunohistochemistry

FFPE sections were deparaffinised with xylene before gradual rehydration and bleached in a 1% KOH/3% H_2_O_2_ solution for 15 minutes. Slides were then stained with haematoxylin, mounted in hydrophilic mounting media and imaged using a Hamamatsu NanoZoomer slide scanner. After coverslip removal, slides were subjected to heat-mediated antigen retrieval in citrate buffer (0.01 M, pH6) for 7 minutes. The sections were then incubated in serum-free protein blocking solution (DAKO) for 30 minutes at room temperature and incubated in primary antibody overnight at 4°C. Following TBS/0.1% tween washes, the sections were incubated in HRP rabbit/mouse secondary antibody (DAKO Real EnVision kit) for 30 minutes at room temperature. Antibody staining was revealed via incubation in AEC chromogen (Abcam) for 5-30 minutes. Following each antibody revealing, the slides were again mounted in hydrophilic mounting media and imaged using a Hamamatsu NanoZoomer slide scanner. The slides were then de-coverslipped and underwent chromogenic de-staining in an alcohol gradient and subsequent antibody stripping via a 75 minute incubation in a solution glycine-SDS, pH 2 at 50 □C, before the next blocking and antibody round (adapted from (Pirici *et al.,* 2009; Tsujikawa et al., 2017)). The following antibodies were used for multiplex IHC: Sox10 (Abcam, ab229331, 1:4,000), mCherry (Abcam, ab125096, 1:4,000) and Mitfa (Sigma, SAB2702433, 1:1,000)

### Statistical analysis

Sample distribution was evaluated using Shapiro-Wilk test. As the samples exhibited non-gaussian distribution, statistical evaluation was performed using nonparametric Wilcoxon test. We set up the threshold to P=0.05 to be considered as statistically significant with the following symbols: ** p<0.01; * p<0.05. Statistical analysis was performed using R software version 4.0.3 through R studio interface version 1.1.456 (R-Core-Team, 2020; RStudio-Team, 2016).

## Acknowledgements

We are grateful to the MRC HGU Zebrafish Facility for zebrafish management and husbandry, Elisabeth Freyer and the IGC Flow Cytometry facility, Ann Wheeler and the IGC Imaging Facility for supporting the imaging experiments. This work was supported by the Cancer Research UK Scotland Centre (CTRQQR-2021\100006). EEP is funded by MRC HGU Programme (MC_UU_00007/9), the European Research Council (ZF-MEL-CHEMBIO-648489), and Melanoma Research Alliance (687306).

## REFERENCES

Avagyan, S., Henninger, J.E., Mannherz, W.P., Mistry, M., Yoon, J., Yang, S., Weber, M.C., Moore, J.L., and Zon, L.I. (2021). Resistance to inflammation underlies enhanced fitness in clonal hematopoiesis. Science 374, 768–772. 10.1126/science.aba9304.

Baron, M., Tagore, M., Hunter, M.V., Kim, I.S., Moncada, R., Yan, Y., Campbell, N.R., White, R.M., and Yanai, I. (2020). The Stress-Like Cancer Cell State Is a Consistent Component of Tumorigenesis. Cell Syst 11, 536–546 e537. 10.1016/j.cels.2020.08.018.

Briona, L.K., Poulain, F.E., Mosimann, C., and Dorsky, R.I. (2015). Wnt/ss-catenin signaling is required for radial glial neurogenesis following spinal cord injury. Dev Biol 403, 15–21. 10.1016/j.ydbio.2015.03.025.

Brombin, A., Simpson, D.J., Travnickova, J., Brunsdon, H., Zeng, Z., Lu, Y., Young, A.I.J., Chandra, T., and Patton, E.E. (2022). Tfap2b specifies an embryonic melanocyte stem cell that retains adult multifate potential. Cell Rep 38, 110234. 10.1016/j.celrep.2021.110234.

Carney, T.J., and Mosimann, C. (2018). Switch and Trace: Recombinase Genetics in Zebrafish. Trends Genet 34, 362–378. 10.1016/j.tig.2018.01.004.

Compton, L.A., Murphy, G.F., and Lian, C.G. (2015). Diagnostic Immunohistochemistry in Cutaneous Neoplasia: An Update. Dermatopathology (Basel) 2, 15–42. 10.1159/000377698.

Dilshat, R., Fock, V., Kenny, C., Gerritsen, I., Lasseur, R.M.J., Travnickova, J., Eichhoff, O.M., Cerny, P., Moller, K., Sigurbjornsdottir, S., et al. (2021). MITF reprograms the extracellular matrix and focal adhesion in melanoma. Elife 10. 10.7554/eLife.63093.

Edelstein, A., Amodaj, N., Hoover, K., Vale, R., and Stuurman, N. (2010). Computer control of microscopes using microManager. Curr Protoc Mol Biol Chapter 14, Unit14 20. 10.1002/0471142727.mb1420s92.

Ellisor, D., and Zervas, M. (2010). Tamoxifen dose response and conditional cell marking: is there control? Mol Cell Neurosci 45, 132–138. 10.1016/j.mcn.2010.06.004.

Ennen, M., Keime, C., Kobi, D., Mengus, G., Lipsker, D., Thibault-Carpentier, C., and Davidson, I. (2015). Single-cell gene expression signatures reveal melanoma cell heterogeneity. Oncogene 34, 3251–3263. 10.1038/onc.2014.262.

Galant, S., Furlan, G., Coolen, M., Dirian, L., Foucher, I., and Bally-Cuif, L. (2016). Embryonic origin and lineage hierarchies of the neural progenitor subtypes building the zebrafish adult midbrain. Dev Biol 420, 120–135. 10.1016/j.ydbio.2016.09.022.

Gemberling, M., Karra, R., Dickson, A.L., and Poss, K.D. (2015). Nrg1 is an injury-induced cardiomyocyte mitogen for the endogenous heart regeneration program in zebrafish. Elife 4. 10.7554/eLife.05871.

Gerber, T., Willscher, E., Loeffler-Wirth, H., Hopp, L., Schadendorf, D., Schartl, M., Anderegg, U., Camp, G., Treutlein, B., Binder, H., and Kunz, M. (2017). Mapping heterogeneity in patient-derived melanoma cultures by single-cell RNA-seq. Oncotarget 8, 846–862. 10.18632/oncotarget.13666.

Jahn, H.M., Kasakow, C.V., Helfer, A., Michely, J., Verkhratsky, A., Maurer, H.H., Scheller, A., and Kirchhoff, F. (2018). Refined protocols of tamoxifen injection for inducible DNA recombination in mouse astroglia. Sci Rep 8, 5913. 10.1038/s41598-018-24085-9.

Johnson, S.L., Nguyen, A.N., and Lister, J.A. (2011). mitfa is required at multiple stages of melanocyte differentiation but not to establish the melanocyte stem cell. Dev Biol 350, 405–413. 10.1016/j.ydbio.2010.12.004.

Jopling, C., Sleep, E., Raya, M., Marti, M., Raya, A., and Izpisua Belmonte, J.C. (2010). Zebrafish heart regeneration occurs by cardiomyocyte dedifferentiation and proliferation. Nature 464, 606–609. 10.1038/nature08899.

Kobayashi, I., Kobayashi-Sun, J., Kim, A.D., Pouget, C., Fujita, N., Suda, T., and Traver, D. (2014). Jam1a-Jam2a interactions regulate haematopoietic stem cell fate through Notch signalling. Nature 512, 319–323. 10.1038/nature13623.

Kwan, K.M., Fujimoto, E., Grabher, C., Mangum, B.D., Hardy, M.E., Campbell, D.S., Parant, J.M., Yost, H.J., Kanki, J.P., and Chien, C.B. (2007). The Tol2kit: a multisite gateway-based construction kit for Tol2 transposon transgenesis constructs. Dev Dyn 236, 3088–3099. 10.1002/dvdy.21343.

Lu, Y., and Patton, E.E. (2022). Long-term non-invasive drug treatments in adult zebrafish that lead to melanoma drug resistance. Dis Model Mech 15. 10.1242/dmm.049401.

Marin-Bejar, O., Rogiers, A., Dewaele, M., Femel, J., Karras, P., Pozniak, J., Bervoets, G., Van Raemdonck, N., Pedri, D., Swings, T., et al. (2021). Evolutionary predictability of genetic versus nongenetic resistance to anticancer drugs in melanoma. Cancer Cell 39, 1135–1149 e1138. 10.1016/j.ccell.2021.05.015.

Marine, J.C., Dawson, S.J., and Dawson, M.A. (2020). Non-genetic mechanisms of therapeutic resistance in cancer. Nat Rev Cancer 20, 743–756. 10.1038/s41568-020-00302-4.

McCloy, R.A., Rogers, S., Caldon, C.E., Lorca, T., Castro, A., and Burgess, A. (2014). Partial inhibition of Cdk1 in G 2 phase overrides the SAC and decouples mitotic events. Cell Cycle 13, 1400–1412. 10.4161/cc.28401.

Mosimann, C., Kaufman, C.K., Li, P., Pugach, E.K., Tamplin, O.J., and Zon, L.I. (2011). Ubiquitous transgene expression and Cre-based recombination driven by the ubiquitin promoter in zebrafish. Development 138, 169–177. 10.1242/dev.059345.

Nguyen, P.D., Gurevich, D.B., Sonntag, C., Hersey, L., Alaei, S., Nim, H.T., Siegel, A., Hall, T.E., Rossello, F.J., Boyd, S.E., et al. (2017). Muscle Stem Cells Undergo Extensive Clonal Drift during Tissue Growth via Meox1-Mediated Induction of G2 Cell-Cycle Arrest. Cell Stem Cell 21, 107–119 e106. 10.1016/j.stem.2017.06.003.

Pan, Y.A., Freundlich, T., Weissman, T.A., Schoppik, D., Wang, X.C., Zimmerman, S., Ciruna, B., Sanes, J.R., Lichtman, J.W., and Schier, A.F. (2013). Zebrabow: multispectral cell labeling for cell tracing and lineage analysis in zebrafish. Development 140, 2835–2846. 10.1242/dev.094631.

Parichy, D.M., Ransom, D.G., Paw, B., Zon, L.I., and Johnson, S.L. (2000). An orthologue of the kit-related gene fms is required for development of neural crest-derived xanthophores and a subpopulation of adult melanocytes in the zebrafish, Danio rerio. Development 127, 3031–3044. 10.1242/dev.127.14.3031.

Patton, E.E., Mueller, K.L., Adams, D.J., Anandasabapathy, N., Aplin, A.E., Bertolotto, C., Bosenberg, M., Ceol, C.J., Burd, C.E., Chi, P., et al. (2021). Melanoma models for the next generation of therapies. Cancer Cell 39, 610–631. 10.1016/j.ccell.2021.01.011.

Patton, E.E., Widlund, H.R., Kutok, J.L., Kopani, K.R., Amatruda, J.F., Murphey, R.D., Berghmans, S., Mayhall, E.A., Traver, D., Fletcher, C.D., et al. (2005). BRAF mutations are sufficient to promote nevi formation and cooperate with p53 in the genesis of melanoma. Curr Biol 15, 249–254. 10.1016/j.cub.2005.01.031.

Pinzon-Olejua, A., Welte, C., Chekuru, A., Bosak, V., Brand, M., Hans, S., and Stuermer, C.A. (2017). Cre-inducible site-specific recombination in zebrafish oligodendrocytes. Dev Dyn 246, 41–49. 10.1002/dvdy.24458.

Pirici, D., Mogoanta, L., Kumar-Singh, S., Pirici, I., Margaritescu, C., Simionescu, C., and Stanescu, R. (2009). Antibody elution method for multiple immunohistochemistry on primary antibodies raised in the same species and of the same subtype. J Histochem Cytochem 57, 567–575. 10.1369/jhc.2009.953240.

R-Core-Team (2020). R: A language and environment for statistical computing. R Foundation for Statistical Computing.

Rambow, F., Marine, J.C., and Goding, C.R. (2019). Melanoma plasticity and phenotypic diversity: therapeutic barriers and opportunities. Genes Dev 33, 1295–1318. 10.1101/gad.329771.119.

Rambow, F., Rogiers, A., Marin-Bejar, O., Aibar, S., Femel, J., Dewaele, M., Karras, P., Brown, D., Chang, Y.H., Debiec-Rychter, M., et al. (2018a). Toward Minimal Residual Disease-Directed Therapy in Melanoma. Cell 174, 843–855 e819. 10.1016/j.cell.2018.06.025.

Rambow, F., Rogiers, A., Marin-Bejar, O., Aibar, S., Femel, J., Dewaele, M., Karras, P., Brown, D., Chang, Y.H., Debiec-Rychter, M., et al. (2018b). Toward Minimal Residual Disease-Directed Therapy in Melanoma. Cell 174, 843–855.e819. 10.1016/j.cell.2018.06.025.

RStudio-Team (2016). RStudio: Integrated Development for R. RStudio, Inc.

Schindelin, J., Arganda-Carreras, I., Frise, E., Kaynig, V., Longair, M., Pietzsch, T., Preibisch, S., Rueden, C., Saalfeld, S., Schmid, B., et al. (2012). Fiji: an open-source platform for biological-image analysis. Nat Methods 9, 676–682. 10.1038/nmeth.2019.

Shen, S., Faouzi, S., Souquere, S., Roy, S., Routier, E., Libenciuc, C., Andre, F., Pierron, G., Scoazec, J.Y., and Robert, C. (2020a). Melanoma Persister Cells Are Tolerant to BRAF/MEK Inhibitors via ACOX1-Mediated Fatty Acid Oxidation. Cell Rep 33, 108421. 10.1016/j.celrep.2020.108421.

Shen, S., Vagner, S., and Robert, C. (2020b). Persistent Cancer Cells: The Deadly Survivors. Cell 183, 860–874. 10.1016/j.cell.2020.10.027.

Singh, A.P., Dinwiddie, A., Mahalwar, P., Schach, U., Linker, C., Irion, U., and Nusslein-Volhard, C. (2016). Pigment Cell Progenitors in Zebrafish Remain Multipotent through Metamorphosis. Dev Cell 38, 316–330. 10.1016/j.devcel.2016.06.020.

Than-Trong, E., Kiani, B., Dray, N., Ortica, S., Simons, B., Rulands, S., Alunni, A., and Bally-Cuif, L. (2020). Lineage hierarchies and stochasticity ensure the long-term maintenance of adult neural stem cells. Sci Adv 6, eaaz5424. 10.1126/sciadv.aaz5424.

Thunemann, M., Schorg, B.F., Feil, S., Lin, Y., Voelkl, J., Golla, M., Vachaviolos, A., Kohlhofer, U., Quintanilla-Martinez, L., Olbrich, M., et al. (2017). Cre/lox-assisted non-invasive in vivo tracking of specific cell populations by positron emission tomography. Nat Commun 8, 444. 10.1038/s41467-017-00482-y.

Tirosh, I., Izar, B., Prakadan, S.M., Wadsworth, M.H., 2nd, Treacy, D., Trombetta, J.J., Rotem, A., Rodman, C., Lian, C., Murphy, G., et al. (2016). Dissecting the multicellular ecosystem of metastatic melanoma by single-cell RNA-seq. Science 352, 189–196. 10.1126/science.aad0501.

Tornini, V.A., Thompson, J.D., Allen, R.L., and Poss, K.D. (2017). Live fate-mapping of joint-associated fibroblasts visualizes expansion of cell contributions during zebrafish fin regeneration. Development 144, 2889–2895. 10.1242/dev.155655.

Travnickova, J., and Patton, E.E. (2021). Deciphering Melanoma Cell States and Plasticity with Zebrafish Models. J Invest Dermatol 141, 1389–1394. 10.1016/j.jid.2020.12.007.

Travnickova, J., Wojciechowska, S., Khamseh, A., Gautier, P., Brown, D.V., Lefevre, T., Brombin, A., Ewing, A., Capper, A., Spitzer, M., et al. (2019). Zebrafish MITF-Low Melanoma Subtype Models Reveal Transcriptional Subclusters and MITF-Independent Residual Disease. Cancer Res 79, 5769–5784. 10.1158/0008-5472.CAN-19-0037.

Tsujikawa, T., Kumar, S., Borkar, R.N., Azimi, V., Thibault, G., Chang, Y.H., Balter, A., Kawashima, R., Choe, G., Sauer, D., et al. (2017). Quantitative Multiplex Immunohistochemistry Reveals Myeloid-Inflamed Tumor-Immune Complexity Associated with Poor Prognosis. Cell Rep 19, 203–217. 10.1016/j.celrep.2017.03.037.

Vendramin, R., Katopodi, V., Cinque, S., Konnova, A., Knezevic, Z., Adnane, S., Verheyden, Y., Karras, P., Demesmaeker, E., Bosisio, F.M., et al. (2021). Activation of the integrated stress response confers vulnerability to mitoribosome-targeting antibiotics in melanoma. J Exp Med 218. 10.1084/jem.20210571.

Wang, L., Wang, S., Yin, J.J., Fu, P.P., and Yu, H. (2009). Light-Induced Toxic Effects of Tamoxifen: A Chemotherapeutic and Chemopreventive Agent. J Photochem Photobiol A Chem 201, 50–56. 10.1016/j.jphotochem.2008.09.013.

Wickham, H. (2016). ggplot2: Elegant Graphics for Data Analysis. Spinger-Verlag.

Wojciechowska, S., van Rooijen, E., Ceol, C., Patton, E.E., and White, R.M. (2016). Generation and analysis of zebrafish melanoma models. Methods Cell Biol 134, 531–549. 10.1016/bs.mcb.2016.03.008.

Wouters, J., Kalender-Atak, Z., Minnoye, L., Spanier, K.I., De Waegeneer, M., Bravo Gonzalez-Blas, C., Mauduit, D., Davie, K., Hulselmans, G., Najem, A., et al. (2020). Robust gene expression programs underlie recurrent cell states and phenotype switching in melanoma. Nat Cell Biol 22, 986–998. 10.1038/s41556-020-0547-3.

Zeng, Z., Johnson, S.L., Lister, J.A., and Patton, E.E. (2015). Temperature-sensitive splicing of mitfa by an intron mutation in zebrafish. Pigment Cell Melanoma Res 28, 229–232. 10.1111/pcmr.12336.

