## Supplementary Figures and Legends for "Fate mapping melanoma persister cells through regression and into recurrent disease in adult zebrafish"

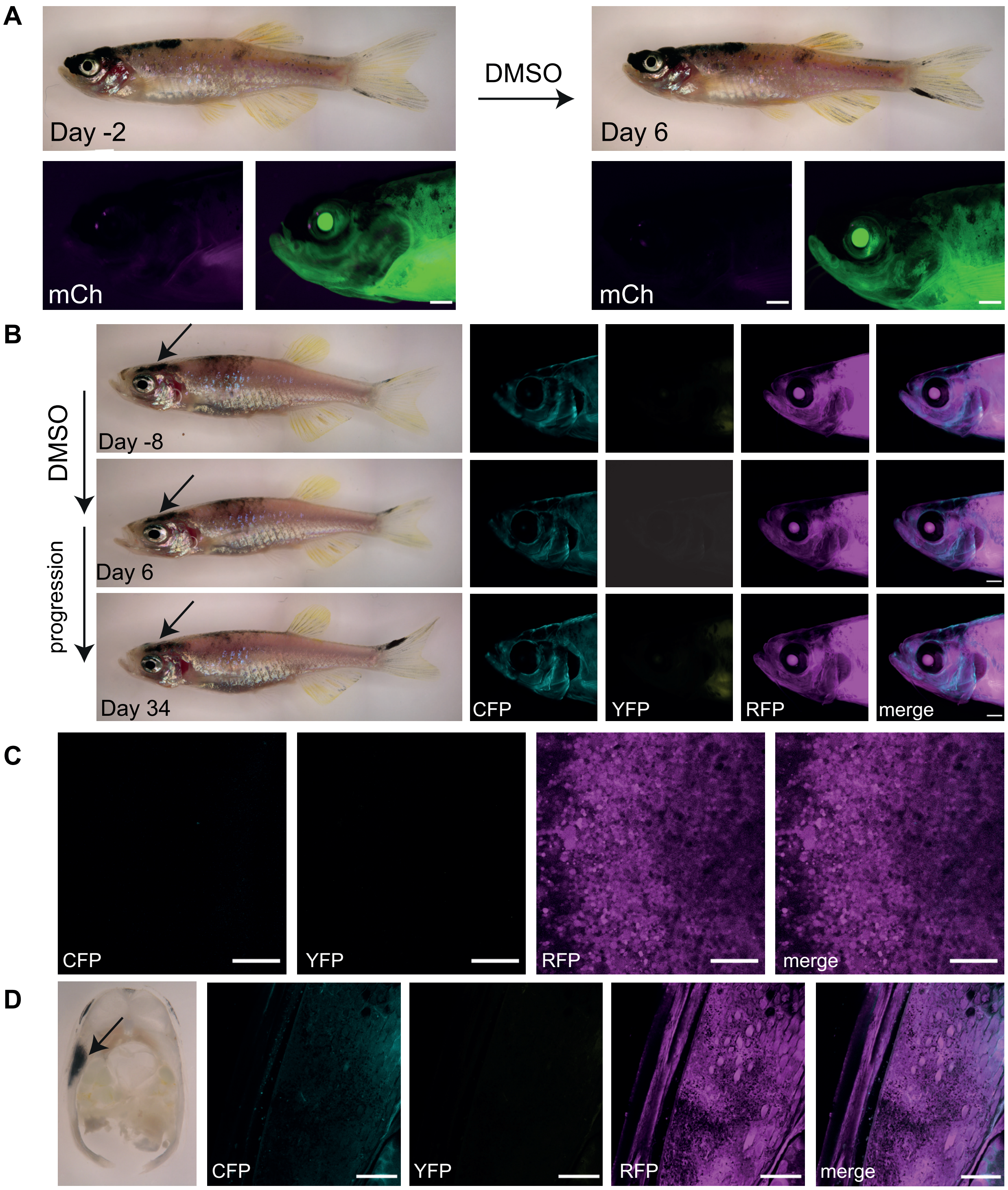


**Figure S1: Control of tamoxifen recombination in the zebrafish melanoma model**

**A.** An adult *mitfa:cre$^{ERt2;}$ubi:Switch* zebrafish with a fully developed primary melanoma tumour 2 days before the DMSO control treatment course (left) and 6 days after the start of the control treatment (right). No de novo expression of mCherry protein is detected in the pigmented head tumour after DMSO control treatment (0.04 %, in magenta). N=2, Scale bar = 1 mm.

**B.** An adult *mitfa:cre$^{ERt2}$;ubi:zebrabow* zebrafish with a primary melanoma tumour 8 days before the DMSO treatment course (top), 6 days (middle) and 34 days after the start of the treatment showing lack of de novo expression of CFP and YFP proteins in tumour after DMSO treatment. N = 3 fish, Scale bar = 1 mm, black arrows point to the tumour location. CFP channel shows some of the non-specific autofluorescence in the creases along jaw and gills independent of tamoxifen induction.

**C.** Single z-plane of confocal acquisition of DMSO-treated control fish (from left CFP, YFP, RFP, merge). In contrast to tamoxifen treated samples in **Figure 3C**, individual cells only express RFP and lack any other fluorophores in the DMSO-treated group. N = 3 fish, scale bar = 100 μm.

**D.** Brightfield image (left) and standard deviation intensity (STD) projections of confocal z-stacks of PFA fixed vibratome section of a DMSO-treated fish. Only RFP expressing cells in the DMSO-treated control group are detected. N = 3 fish, scale bar = 200 µm, black arrow points at tumour location.

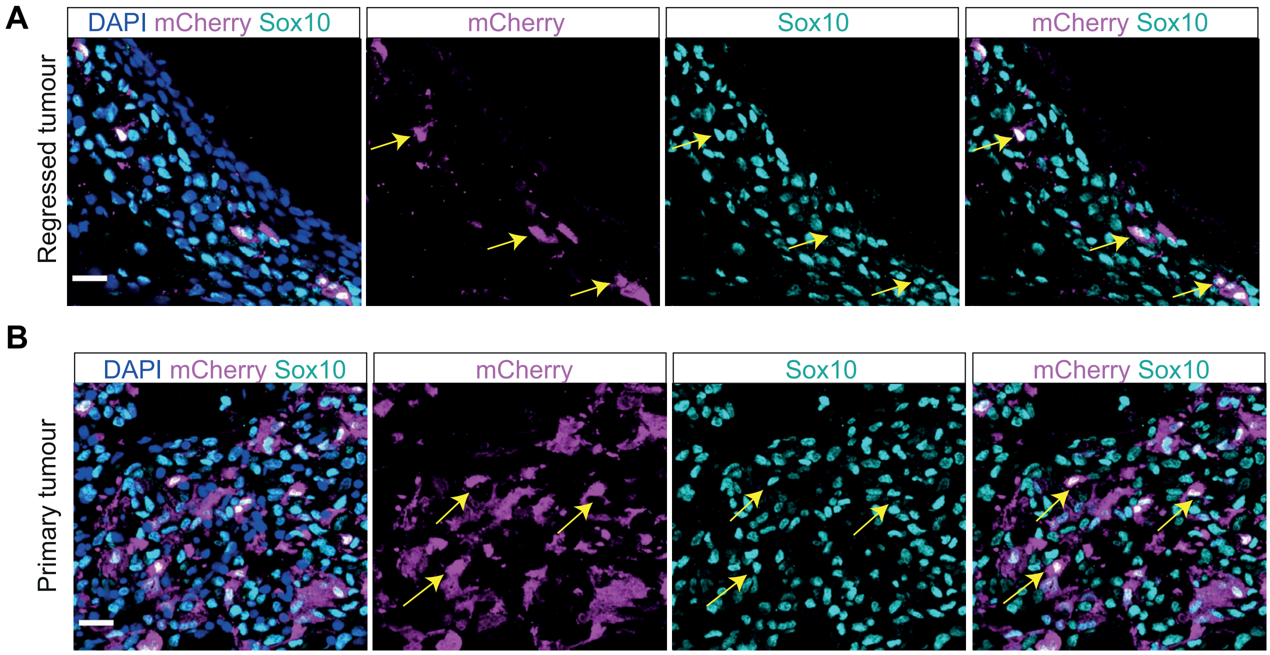


**Figure S2: Immunofluorescence validates that the mCherry positive cells in the regressed tumour co-express the melanoma marker Sox10.**

**A.** STD projections of confocal z-stack acquisitions of immunofluorescence staining of the tamoxifen-treated regressed tumour, showing staining of mCherry and Sox10 proteins with DAPI nuclear staining. Yellow arrows indicate mCherry positive tumour cells that are also positive for Sox10. Scale bar = 15 μm.

**B.** STD projections of confocal z-stack acquisitions of immunofluorescence staining of the tamoxifen-treated primary tumour, showing staining of mCherry and Sox10 proteins with DAPI nuclear staining (yellow arrows). Scale bar = 15 μm.

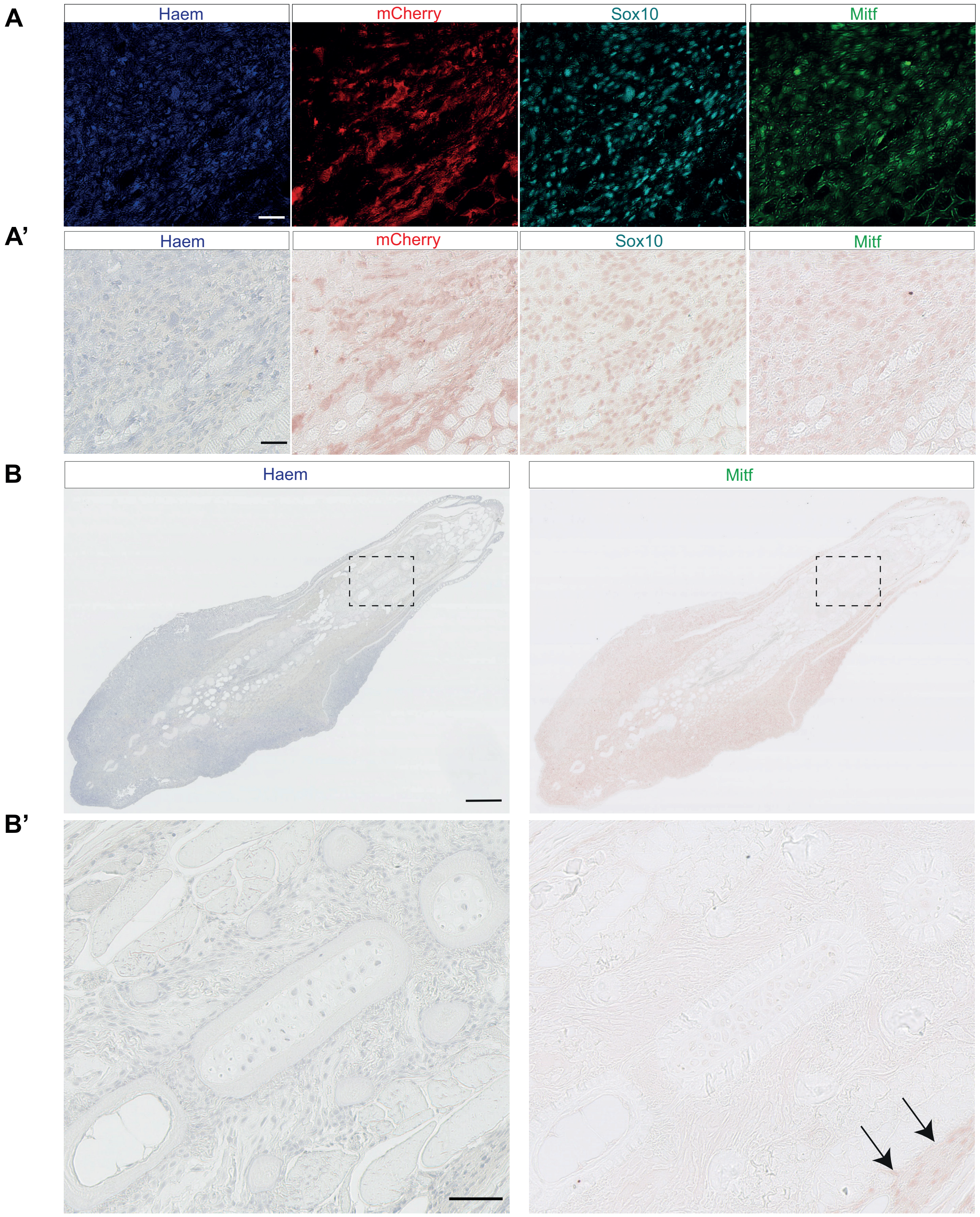


**Figure S3: Matched brightfield and pseudo-coloured images of tamoxifen treated recurred tumour sections.**

**A.** Enlarged views of the tumour section shown in Figure 7D, showing haematoxylin nuclear counter-stain and individual staining of Sox10, Mitfa, and mCherry proteins. Scale bar, 25 µm.

**A’.** Corresponding bright field images to A, prior to pseudo-colouring on ImageJ. Scale bar, 25 µm.

**B.** Overview of the brightfield images of the recurred tumour section shown in Figure 7D, showing haematoxylin nuclear counterstaining and staining of Mitfa protein. Scale bar, 250 µm. Grey dashed boxes indicate areas of section shown in B’.

**B’.** Enlarged views of brightfield tumour sections shown in B, showing lack of Mitfa staining in non-tumour cells which are positive for haematoxylin and are adjacent to the tumour. The edge of tumour tissue that is positive for Mitfa staining is visible (black arrows). Scale bar, 25 µm.
